# Criteria for evaluating molecular markers: Comprehensive quality metrics to improve marker-assisted selection

**DOI:** 10.1101/249987

**Authors:** John Damien Platten, Joshua N. Cobb, Rochelle E. Zantua

## Abstract

Despite strong interest over many years, the usage of quantitative trait loci in plant breeding has often failed to live up to expectations. A key weak point in the utilisation of QTLs is the “quality” of markers used during marker-assisted selection (MAS): unreliable markers result in variable outcomes, leading to a perception that MAS products fail to achieve reliable improvement. Most reports of markers used for MAS focus on markers derived from the mapping population. There are very few studies that examine the reliability of these markers in other genetic backgrounds, and critically, no metrics exist to describe and quantify this reliability. To improve the MAS process, this work proposes five core metrics that fully describe the reliability of a marker. These metrics give a comprehensive and quantitative measure of the ability of a marker to correctly classify germplasm as QTL[+]/[-], particularly against a background of high allelic diversity. Markers that score well on these metrics will have far higher reliability in breeding, and deficiencies in specific metrics give information on circumstances under which a marker may not be reliable. The metrics are applicable across different marker types and platforms, allowing an objective comparison of the performance of different markers irrespective of the platform. Evaluating markers using these metrics demonstrates that trait-specific markers consistently out-perform markers designed for other purposes. These metrics also provide a superb set of criteria for designing superior marker systems for a target QTL, enabling the selection of an optimal marker set before committing to design.

## Introduction

The world population is expected to top 9 billion people by 2050. To feed this population, it is estimated that agricultural output of cereals alone will need to increase by approximately 1 billion tons [1]. It is widely acknowledged that meeting this growth target will require the integration of new technologies into the breeding process. Many authors have discussed the promise of molecular marker technologies for improving the speed and efficiency of the breeding process, and extensive literature has accumulated on methodologies to incorporate the use of markers into breeding decisions (e.g. [2]). At its core, this marker-assisted selection (MAS) is about two correlations:

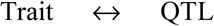

and

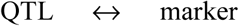

A QTL is identified as a genetic position (locus) associated with some degree of phenotypic variation in a specific trait. Markers are assayable polymorphisms with some degree of association with a QTL in a specific gene pool. Both sets of correlations may be broad-ranging or narrowly applicable, and the success or reliability of MAS is directly determined by the strength of these correlations. However, since the middle factor (QTL) is almost never tracked *per se*, both correlations are often conflated into the indirect association of the marker with the trait. Since this association is indirect, reliable markers may not always result in reliable improvement of the trait – this depends on the quality of the QTL, and is a topic for a different discussion. However, unreliable markers will always result in unreliable improvement, and it is thus essential to identify what constitutes a reliable marker, and metrics to objectively measure and evaluate this quality.

Current MAS programs use one of several genotyping platforms, depending on requirements for marker density and sample throughput. These platforms range from low-throughput, PCR-based techniques such as the traditional microsatellite/simple-sequence repeats (SSRs) and newer insertion/deletion mutations (indels), to the explosion of high-throughput single-nucleotide polymorphism (SNP) platforms and new sequencing-based methods such as genotyping-by-sequencing (GBS) and amplicon sequencing [3, 4, 5, 6]. Most recent literature has focused on new SNP technologies, but by far the most common systems in use by public sector breeding programs are traditional SSRs. These are widely employed in biparental mapping studies such as QTL mapping and fine-mapping [7, 8, 9, 10, 11, 12, 13, 14], but have also been used in cross-population meta-analyses [12] and allelic diversity assessments [15, 16]. Given their high throughput nature SNPs and GBS are the platforms of choice for strategies requiring high sample volumes and/or marker densities, such as genomic selection and genome-wide association studies (e.g. [17, 18]).

Despite the acknowledged importance of incorporating molecular markers into the breeding process, there has been little discussion on factors influencing the success of such endeavours. Some studies have investigated the utility of SSRs in breeding processes [15, 19, 20, 21], however the only criterion for usefulness that is considered is genetic linkage with the QTL; other issues such as how reliably the markers classify favourable and unfavourable alleles are rarely examined, and if so done in a cursory manner.

Poor classification ability in markers can lead to many undesirable outcomes. For example, a typical MAS workflow involving SSRs starts with a parental polymorphism survey; SSRs are chosen to introgress a QTL based on their linkage (position) and the fact they are polymorphic between a donor and the chosen recipient line. However, this does not examine whether the chosen recipient already contains the target QTL; the ability of the QTL to improve the recipient is assumed, not tested. SSR markers cannot provide information on whether the chosen recipient already possesses the QTL - this is a circular argument - and there are documented cases where SSRs give misleading indications as to the presence/absence of a QTL. This can be seen for example in [15] and [22], which in both cases confuse varieties with different *Saltol* alleles and distinguish varieties with the same allele (compare for example [23]). Many similar situations have arisen in breeding programs, irrespective of marker platform in use (SNP or gel-based), leading to unreliable outcomes, usually as a result of classifying a variety as lacking a QTL when in fact the QTL is already present. To avoid similar problems in future, some method of characterising markers is required, to help identify markers at risk of giving misleading results and to aid in the design of superior markers as replacements. Most importantly, a set of objective measures should be derived that describe how accurately a marker classifies both QTL[+] and QTL[-] material.

Surprisingly there is no literature available dealing with this subject, which may contribute to the ambiguity surrounding the quality of existing marker systems. The closest parallels in other disciplines are accuracy metrics used to evaluate clinical tests (e.g. [24]), but the concepts do not map directly to each other. For example, medical diagnostics assume a large number of case-[+] and case-[-] datapoints are available, and often deal with quantitative measures such as enzyme or antibody activity levels. By contrast, “datapoints” in the case of molecular markers are characterised varieties with or without the target QTL, which are typically far fewer in number. Combined with this, for many genotyping platforms the genotype is essentially a binary output, making it strictly impossible to distinguish more than two allelic states.

To stimulate discussion in this area, a set of five core and nine supporting metrics is presented along with strategies for their calculation, which attempt to capture the level and type of association between a marker and its target QTL. These metrics are focused primarily on assessing the classification ability of a marker against a background of high allelic polymorphism such as is found in rice [25], but also assess several other parameters related to the reliability of scoring and usefulness of a marker in breeding. The metrics are then used to evaluate a set of SSR, SNP and indel markers targeting a range of QTLs in rice (*Oryza sativa* L.), to illustrate their application in designing marker systems that give higher confidence for deployment in breeding programs.

## Materials and Methods

A summary of proposed marker quality metrics is found in Table 1. The metrics are comprised of five core metrics that quantify the reliability of a marker, and a further nine supporting metrics that allow the determination of the core metrics. They fall into three main categories: (1) Technical metrics, (2) Biological metrics, and (3) Breeding metrics. To evaluate these metrics on existing marker systems in rice several different kinds of markers from multiple platforms were compared. These include 20 SSR markers [3], approximately 4500 SNP markers identified as part of the OryzaSNP project [25] and utilised on fixed genotyping platforms such as the Illumina Infinium chip ([6]; hereinafter the “anonymous SNP panel”), and 137 candidate QTL-specific SNP and indel markers. A list of markers evaluated can be found in S1 Table. Markers were evaluated using one of two main datasets, depending on the metrics to be evaluated: empirically-determined PCR results to compare technical performance and breeding metrics for gel-based SSR and indel markers, and a large genome resequencing dataset to compare biological and breeding metrics for anonymous SNPs, QTL-specific SNPs, and QTL-specific indel markers.

**Table 1.** Summary of core and supporting metrics to describe marker quality

| Technical metrics |  |  |
| --- | --- | --- |
| Version | Support | Identifier designating the version of a marker being examined; particularly important for technical performance metrics. |
| Call rate | Core | The proportion of samples which give a scorable result. For gel-based markers this is easy to visualise and understand (unless the marker is dominant), but many SNP platforms will call an allele even if the underlying data is substandard. |
| Clarity | Core | How reliably a sample can be classified as allele A, B or heterozygous. This is a critical parameter, and for each new marker it is worthwhile determining – even commercial SNP platforms do not always perform to the standard the sales brochure will advertise, and each individual marker will perform in different ways. The clarity can be quantified by determining what proportion of known germplasm is correctly classified, and/or how well it correlates with tightly-linked alternative markers. |
| Biological metrics |  |  |
| Linkage | Support | Genetic distance of the marker from the QTL peak, it is probably best expressed as a genetic distance gained from one or more mapping populations. Diagnostic markers will by definition have a distance of 0cM. |
| Position | Support | Position targeted by a marker, for example chromosomal location (preferable), mapping bin or consensus genetic position. |
| Derived QTL state | Support | Which state appears to be the most derived, and therefore most informative in determining presence of the QTL: Favourable or Unfavourable. This is not a property of a specific marker, but rather the QTL. |
| Marker Target | Support | A qualitative description of the allele targeted by the marker, the favourable or unfavourable allele. |
| Favourable allele | Support | Not quality metrics per se but necessary for automated analyses, and helpful for new researchers wanting to use a marker. Depending on the platform these could be a size (143bp, Large/Small), allele (A/C/G/T) or even an allele definition (e.g. for amplicon sequencing). |
| Unfavourable allele |  |  |
| False Positive Rate | Core | The proportion of known negative genotypes incorrectly classified as QTL[+]. Assayed as the number of *known* recipients identified as not having an unfavourable allele(s) of the marker (and thus incorrectly classified as QTL[+]).<br>$\frac{(\# \text{ recipients withOUT unfavourable alleles})}{(\text{Total } \# \text{ known recipients})}$ |
| False Negative Rate | Core | This is the converse of FPR, i.e. the proportion of known QTL[+] genotypes incorrectly classified as QTL[-] due to not having a favourable allele of the marker.<br>$\frac{(\# \text{ donors withOUT favourable alleles})}{(\text{Total } \# \text{ known donors})}$ |
| Breeding metrics |  |  |
| Breeding program False Positive Rate | Support | Analogous to the FPR and FNR respectively, but assessed on a particular breeding panel. Since the breeding panel likely has a lower level of allelic diversity, markers should score higher on the BpFPR and BpFNR than on the true FPR/FNR. These metrics give more precise information on the behaviour of a marker in the target breeding program, but the association is then specific to that program, so must be reassessed for other programs. |
| Breeding program False Negative Rate |  |  |
| Utility | Core | The proportion (percentage) of a prospective breeding pool across which a marker could be used to introgress a QTL. This is equivalent to the proportion of the pool which does NOT carry the donor allele of a marker:<br>$\frac{(\# \text{ cultivars withOUT favourable alleles})}{(\text{Total } \# \text{ cultivars assessed})}$ |

### Technical metrics: Call rate and clarity

Technical metrics such as call rate and clarity must be determined empirically from genotyping data using a specific marker assay (primers, probes, etc.); it is entirely possible for two independent sets of primers targeting the same locus to give vastly different results on these metrics. To this end, a set of 20 SSR and 86 trait-specific indel markers (S1 Table) were empirically evaluated on a set of 121 diverse varieties released by the International Rice Research Institute and others, supplemented with various QTL donor and recipient germplasm characterised as part of QTL mapping exercises by numerous groups. A list of varieties examined is found in S2 Table. Indel marker genotypes were scored as Large/Small, while SSR products were assigned to band size categories as appropriate. Missing and unclear results were flagged as such. The call rate was determined as the percentage of samples giving a visible result (i.e. not “missing”), while clarity was the percentage of samples giving a clearly scorable result (i.e. as opposed to unclear or ambiguous results).

### Biological and breeding metrics: false positive rate, false negative rate, and utility

These metrics need to be determined against a background of high allelic diversity. To achieve this, whole-genome resequencing data was obtained for a set of 242 diverse rice accessions, comprising 173 cultivated lines (named, released varieties) and 69 landraces, most of which were chosen for their status as QTL donors or recipients. Much of this data was obtained from the rice 3000 genome dataset [26], supplemented with resequencing of high-value donors and recipients for specific QTLs. A list of varieties examined is found in S3 Table. Raw data (reads) were mapped to the MSU7 rice genome build using bwa, and resulting bam files processed using samtools [27, 28]. A total of 352 anonymous SNPs (i.e. not designed specifically for a given QTL) were chosen from the ∼4500 useable features represented on the Infinium SNP chip [6], either within QTL limits (for large QTLs) or within similar distances to the QTL-specific SNP and indel markers. A total of 482 QTL-specific markers, both SNP and indel, were chosen within QTL limits, or within short physical distances of known, cloned genes. Details of the marker positions examined are found in S4 Table. Nucleotide base calls were obtained at all SNP sites – both QTL-specific SNPs and anonymous SNP markers – using standard Samtools/bcftools pipelines [27]. Genotype calls for QTL-specific indel marker positions were determined manually from the same dataset, as automated variant-calling algorithms did not produce reliable results for indels >∼5nt.

All data was consolidated in a MS Access database and information on favourable and unfavourable alleles was recorded for each marker. For anonymous SNPs, which do not have defined favourable or unfavourable alleles, data on accuracy metrics was calculated in two ways: first using the assigned allele A as favourable and B as unfavourable, and secondly classifying the allele with the highest frequency in known donor lines (lowest false negative rate; FNR) as favourable. The latter method was designated as “FNR corrected”. Data was analysed across 42 target QTLs for various disease resistance, abiotic stress, yield and flowering-related traits (S5 Table). As each QTL typically spanned a significant physical distance, summary data was calculated first across all markers of a particular type within a QTL, then averaged across the 42 target QTLs. The final dataset consisted of 352 anonymous SNPs, 251 trait-specific SNPs and 223 trait-specific indels, scored across a common set of 242 genomes for the 42 target QTLs.

## Results

### Derivation of quality metrics

In developing a set of metrics to assess the performance of a candidate marker, it is necessary to break down the features of a marker that impact on its reliability. Broadly speaking, markers may vary in three main areas:

1. Technical aspects related to the assay and scoring of the marker;
2. Biological aspects of the association between the marker and its target locus;
3. Practical aspects of the marker’s use in a breeding pool.

These three areas are quite independent of each other; there are many markers which score well in some categories but fail completely in others, and thus for an accurate picture of how a marker behaves, all three areas must be examined. In addition, to adequately characterise a marker in each area, the areas themselves must be broken down into several measurable quality metrics (Table 1). The metrics are divided into five core metrics that quantify the reliability of a marker, and another nine supporting metrics that enable the calculation of the core metrics and deal with more difficult case studies (Fig 1).

**Fig 1.**
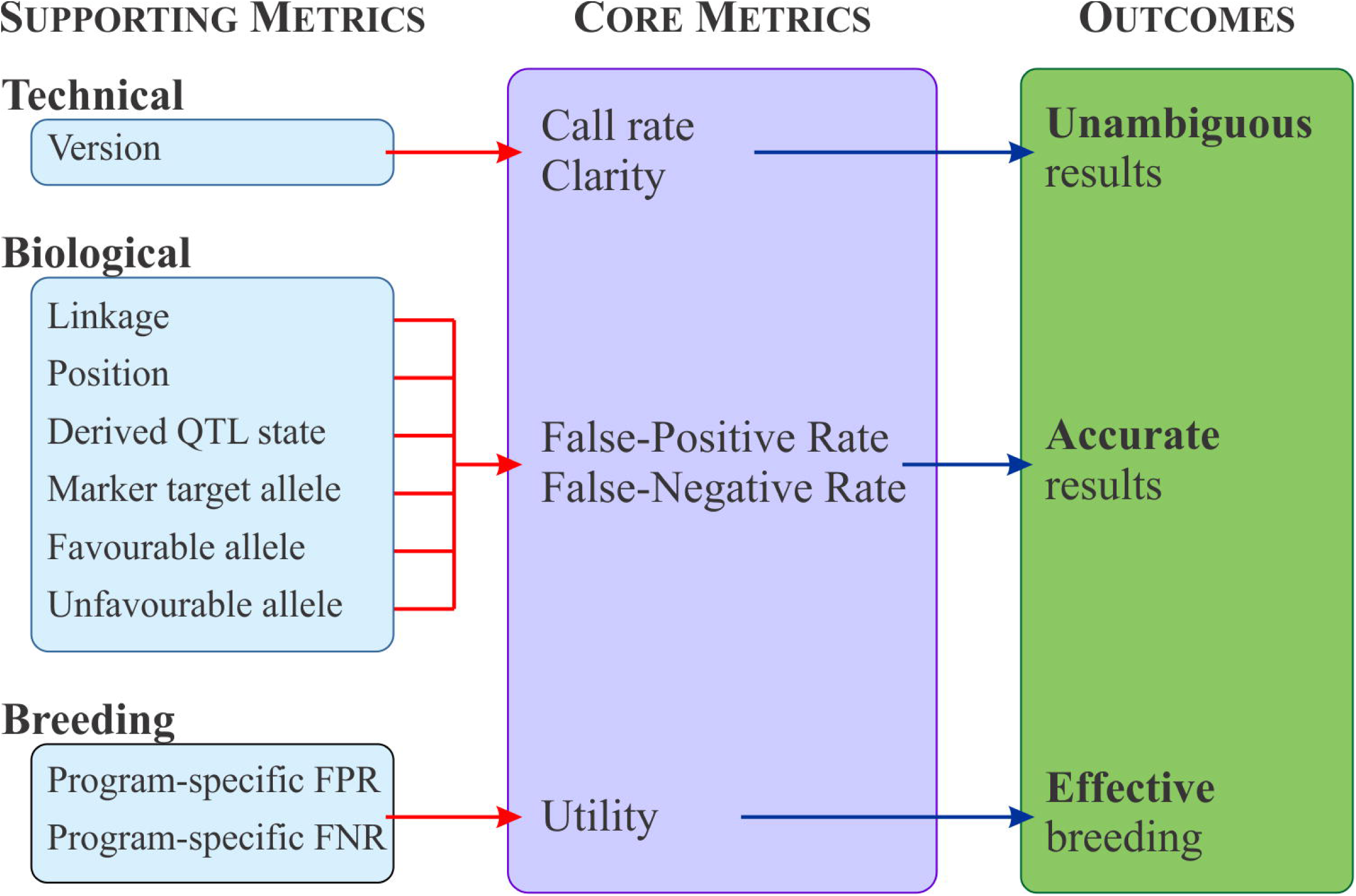
Overview of the marker quality metrics. Core metrics capture the most critical information relating to marker performance, accuracy and usefulness. Supporting metrics are needed in the calculation of the core metrics and/or capture other important information, but are not routinely required in making breeding decisions.

### Technical metrics

Technical metrics relate to how confidently a randomly selected sample from a genotyping job gives an accurate result and can be captured clearly by two core metrics: call rate and clarity. Call rate is the proportion of samples that give a scorable result (as opposed to a “missing” result). Many commercial genotyping platforms already report call rates as a metric of platform performance; typical claims are >99%. Less often reported are estimates of a marker’s clarity. At its simplest, clarity is a subjective opinion on how *clear* the results are, i.e. how reliably genotypes A, B and H can be distinguished. In a more objective sense, an estimate of this could be obtained from how often samples with known genotype are reported to have the correct score, or how often duplicate samples match. Commercial SNP platforms occasionally report statistics on clarity (or repeatability), but without recourse to raw data – which is rarely available – these are difficult to verify. Finally, since every existing technology makes some use of target-specific oligonucleotides, different marker assays on the same platform will vary in their quality on these metrics, even if they target the same position and polymorphism. Thus, versioning of the marker is necessary to allow distinguishing the performance of alternate forms of a marker.

### Biological metrics

Biological metrics can be broken down into qualitative metrics that describe the *type* of association between a marker and QTL, as well as quantitative metrics that describe the *level* of association. These are the most important metrics for successful MAS, and also the most complex. The root cause of differences in marker associations stem from the evolution of traits in an organism, and specifically the relative evolutionary timelines in which the causative allele for a QTL and the polymorphism at the proposed candidate marker emerged. An illustration of this is given in Figs 2 and 3. In all cases, irrespective of whether the causal allele is favourable or unfavourable, the polymorphism most reliable in classifying the donor and recipient lines is one which arose at the same time (and in the same lineage) as the mutation giving rise to the causal, derived allele. This is because the causal mutation and the marker allele are in perfect LD when they emerged and remain in perfect LD in the gene pool consistent with the probability of a recombination event between them.

**Fig 2.**
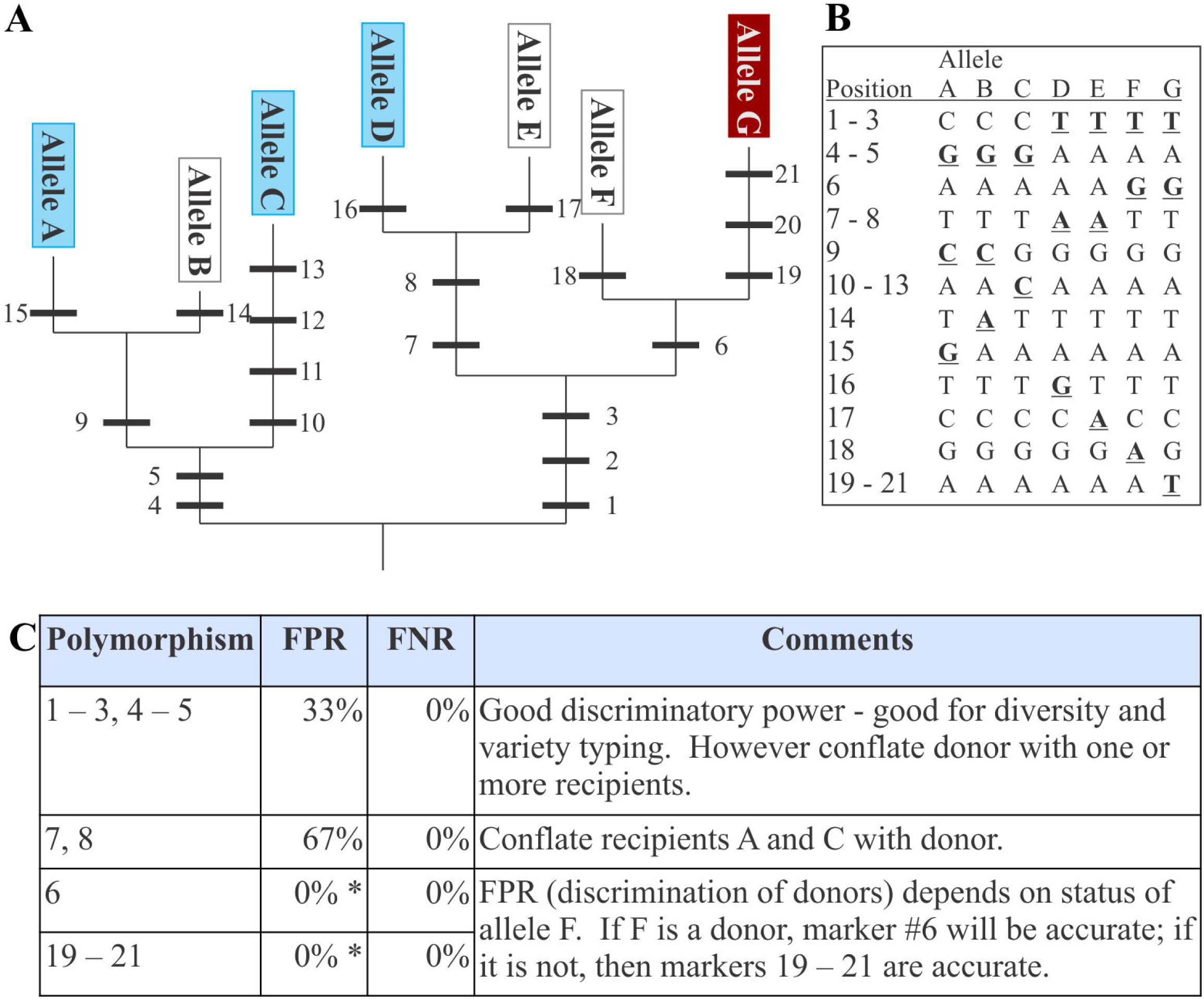
Illustration of the evolution of a QTL. **A** Starting from an ancestral point, mutations in a particular gene accumulate (numbered black bars, representing mutations in **B**), resulting in new alleles. At some point, a mutation arises which improves a trait, resulting in a donor allele for a QTL (dark/red), distinguishing it from known recipient alleles (light/blue); typically the status of many alleles is unknown (white). **C** Each mutation (1 – 21) is a potential marker, and all are found in the same gene, but some are more informative than the others. Comparing the false positive and false negative rates for each mutation allows the determination of which polymorphism gives the most reliable discrimination between the donor and recipient phenotype classes.

**Fig 3.**
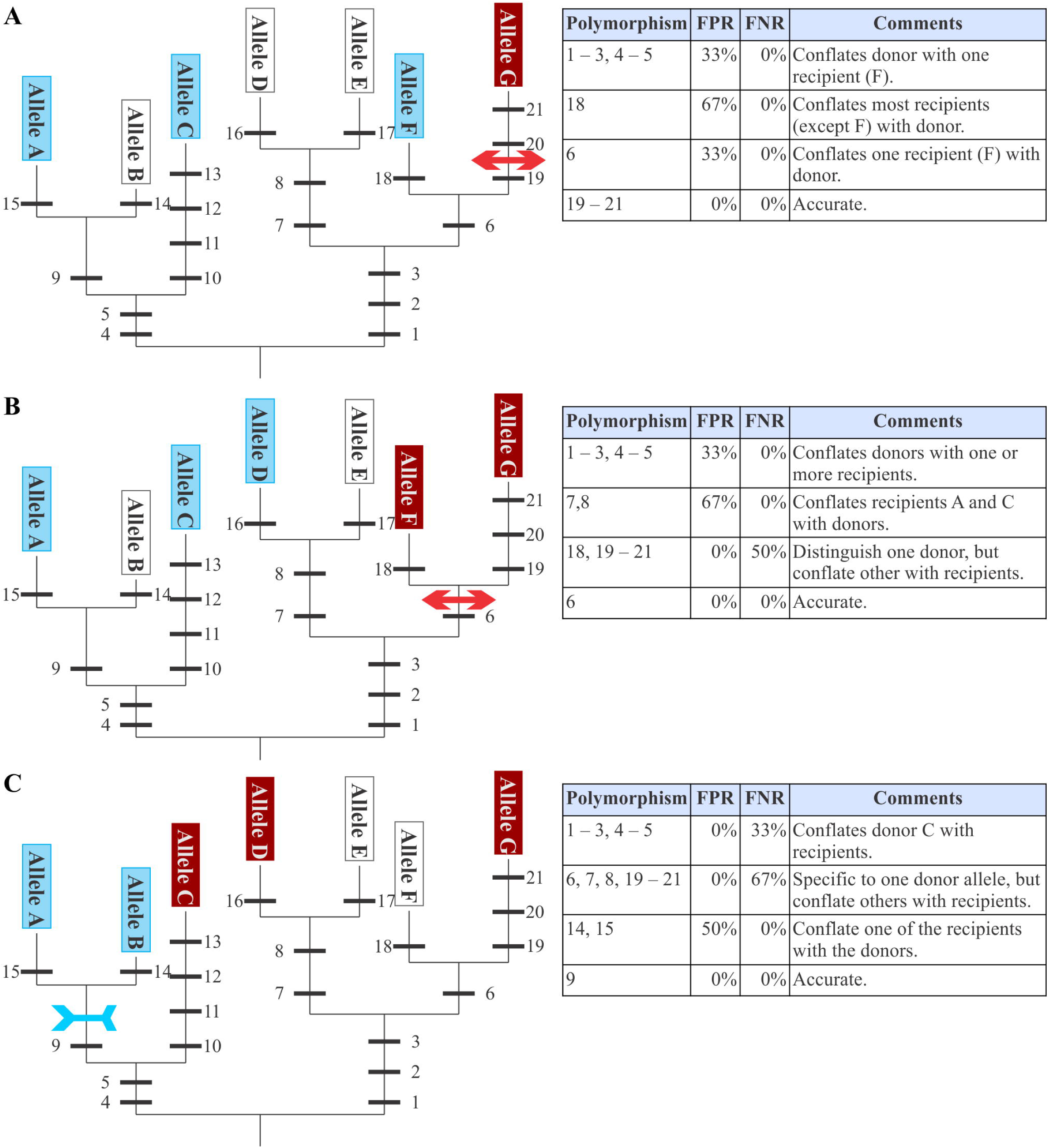
Determining the optimal polymorphism under several evolutionary scenarios. Depending on when the causative mutation arose (arrowed bars; outwards pointing for mutations conferring favourable or inwards facing for unfavourable alleles respectively), there may be only one favourable allele (**A**), a small number of alternative favourable alleles and multiple unfavourable alleles (**B**), or multiple favourable alleles and a few unfavourable (**C**). In **A** and **B**, the derived allele for the QTL is the favourable allele(s); in **C** it is the unfavourable. In all cases the polymorphism which gives the most accurate classification of donor and recipient status is one which arose in the same lineage and at a similar time to the causal, derived mutation.

From this theoretical consideration, it is clear that a number of parameters must be specified in order to accurately describe the association of a marker with its target QTL. Descriptive metrics such as which allele of the QTL (favourable or unfavourable) represents the derived state, the allele (favourable or unfavourable) targeted by a marker, and specifications of marker linkage, favourable and unfavourable alleles, all describe the *type* of association the marker has with its target QTL. Most are easily determined, although determining which QTL allele is derived may require a detailed genomic investigation. Nonetheless, if this can be determined accurately, then markers specific to the derived allele have the greatest chance of also providing a reliable classification across novel allelic diversity, reducing the risk of incorrect classification in future breeding efforts. Therefore, expending some effort to determine the derived QTL allele before designing large numbers of markers is justified.

The proposed quantitative QC metrics of false positive rate and false negative rate (FPR and FNR) describe the *level* of association between the marker and its target QTL and are arguably the most important of the metrics presented, but also the most difficult to estimate. The FNR is the proportion of known donor lines that are incorrectly classified as QTL[-] by the marker. Since the lines are known to carry favourable alleles of the QTL, classification of any of these lines as QTL[-] thus represents a false-negative call by the marker. This is the converse of the marker’s specificity. Markers with a high FNR pose a significant risk of mis-classifying samples as QTL[-] when they do in fact carry a favourable allele; thus breeding material may be discarded which in reality could have been advanced. Markers with a low FNR will correctly identify all samples that possess the QTL[+] state but may still mis-classify samples with an unfavourable (non-donor) alleles as QTL[+], in other words a low FNR does not imply a low FPR.

The FPR is simply the converse and represents the proportion of non-donor (recipient) lines that are incorrectly classified as QTL[+], and is the converse of the marker’s sensitivity. In a breeding context, a high FPR means there is a significant risk of investing in and advancing lines based on MAS results that indicate the presence of the QTL[+] allele, but in reality are QTL[-].

It is important to reiterate here that the FPR and FNR are the proportion of lines with *known* QTL[+]/[-] alleles that are correctly classified as such. They are fundamentally linked to the diversity of alleles with known function, which is determined by the effort that has been put into characterising/mapping donor and non-donor diversity. Many markers are chosen for breeding applications based on their linkage with QTL alleles in specific mapping populations where the QTL is discovered. But those markers – while informative in the chosen mapping population – could still score poorly on both FPR and FNR, resulting in poor performance once deployed as MAS targets. The difficulty arises because the marker is being applied to new populations, where at least one of the parents is of unknown status with respect to the QTL. Markers with low FPR and low FNR will faithfully report the presence or absence of the QTL in any sample, irrespective of allelic diversity in any gene pool of interest.

### Breeding metrics

Breeding QC metrics describe the relative value of applying a marker in a specific breeding program. These consist of three metrics: Breeding program false positive rate (BpFPR), Breeding program false negative rate (BpFNR) and Utility. BpFPR and BpFNR are equivalent to the FPR and FNR metrics described above, but are specific to particular breeding program in which they are assessed, rather than on the full diversity of known donors and recipients. Since the breeding pool may be expected to have lower allelic diversity than occurs species-wide, and because selection and genetic drift are modifying patterns of LD independently across breeding programs, these rates can be quite different from the true FPR and FNR (the usual expectation would be the breeding program rates to be lower than the true rates, though the opposite could also occur). They may also be different for different breeding programs, and must be assessed independently for each. They will require the determination of donor and recipient lines *within a breeding program*, which will involve collecting phenotype data for each program under investigation. But once gathered, the QC metrics directly quantify the marker’s reliability for making breeding decisions in that specific program. It’s worth noting however, that this is predicated on the assumption that the assessed panel represents the complete allelic constituency present in a breeding program, and that the breeding strategy focuses on increasing the frequency of favourable haplotypes through recombination in a closed gene pool, minimizing the introduction of novel allelic variation which may introduce marker alleles that are not in LD with the causal polymorphism.

Finally, utility is the proportion of a breeding panel over which a marker could be used to select for its associated QTL[+] allele. This is basically an assessment of the frequency of QTL[+] alleles in the breeding program, easily calculated as the proportion of breeding lines which possess *non-donor* allele(s) of the marker (Fig 4). A program where the marker offers high utility by definition has the QTL[+] allele at low frequency. Note that this is entirely separate from the FPR and FNR: a marker may perfectly classify all material as to its QTL status, but if the QTL is fixed in the breeding program then the marker (QTL) is of little utility in improving the trait. A good example of this in rice would be *sd1*, which due to intense selection pressure for plant height and heavy usage of QTL[+] *sd1* green revolution varieties by breeding programs is fixed in nearly all breeding populations and therefore is not available to manipulate plant height.

**Fig 4.**
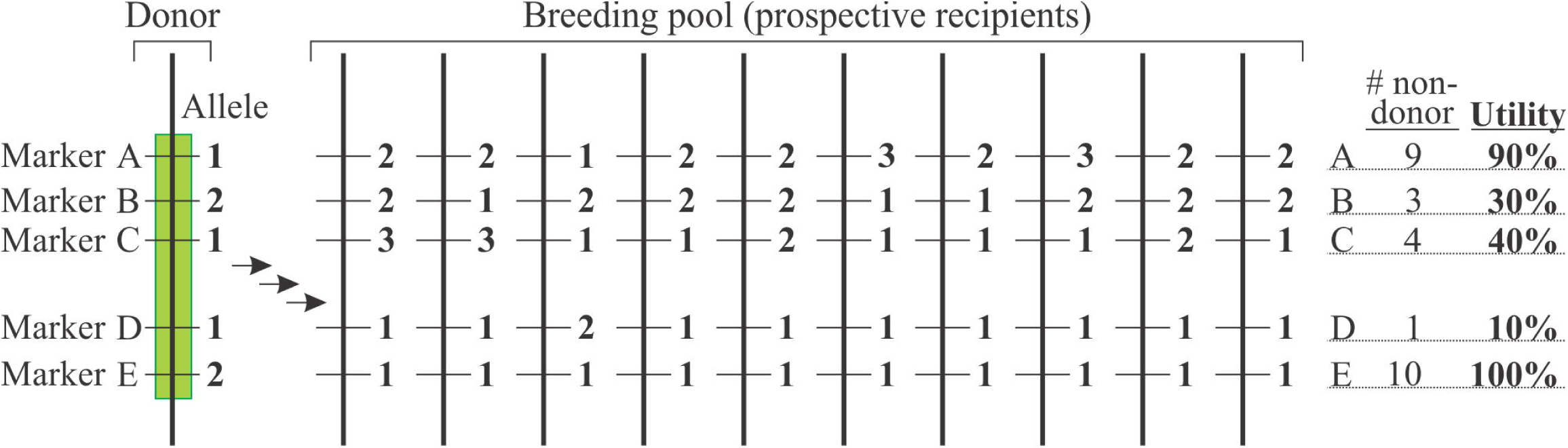
Utility of a marker. Alternative markers within a QTL region (markers A – E) each have multiple alleles (numbered). Alternative alleles of each marker are found at differing frequencies within a breeding pool. Those markers with a high frequency of the favourable allele in the breeding pool (B, C, D) – and thus low utility – can only distinguish the donor genotype in a small number of breeding backgrounds. By contrast markers A and E have high or very high utility, as they are polymorphic with respect to nearly all target genomes in the breeding pool.

A summary of all the proposed metrics is presented in Table 1. Several of these metrics are purely descriptive (derived QTL allele, marker target allele, donor and recipient alleles), but are required to allow the calculation of the more quantitative parameters; these are called supporting metrics. The quantitative parameters then provide a detailed assessment of the performance of a marker; these are the core metrics, and provide the best criteria for assessing markers. Ideal target values and consequences using markers with unfavourable performance scores using these metrics are explained in Table 2. Of particular note are the core metrics FPR and FNR; poor scores on these will increase the probability of discarding valuable breeding germplasm, or worse, wasting resources advancing QTL[-] lines. Assuming good scores on FPR and FNR (or BpFPR and BpFNR), a poor utility value indicates the marker (and thus the QTL[+]) allele is nearly monomorphic in the breeding program and can only be used to select for the QTL across a narrow/small proportion of the breeding pool. Finally the derived marker allele (donor/recipient) should ideally match the derived allele of the QTL; if so, the marker stands a much better chance of correctly classifying additional unknown or uncharacterised alleles.

**Table 2.** Ideal target values for key metrics

| Metric | Target value | Consequences of unfavourable score |
| --- | --- | --- |
| Call rate | 100% | High number of samples with missing data (cannot be genotyped); reduction in efficiency of program and possibility of missing high-value germplasm. |
| Clarity | 100% | High number of samples with ambiguous genotype. Possibility of misclassifying material in the donor/recipient categories; propagating germplasm with low value, or discarding germplasm with high value. |
| Linkage | 0cM | High chance of recombination between the marker and actual QTL, leading to a breakdown in the marker-trait association. |
| Marker target | Derived QTL state | Misclassification of new, unrecognised or uncharacterised allelic diversity. This is of particular use in situations where few donor and recipient lines have been characterised; if the marker target allele matches the derived QTL allele (favourable or unfavourable), the marker is more likely to correctly classify new, uncharacterised alleles. |
| FPR/BpFPR | 0% | Failure to distinguish some unfavourable alleles from some favourable alleles; reports presence of QTL when it is actually absent. May result in use of a line as donor when it is not, or a lack of effort to use a QTL to improve a trait when this would be appropriate. This is in general a more serious failure than low FNR. |
| FNR/BpFNR | 0% | Failure to distinguish some favourable alleles from unfavourable ones; reports absence of QTL when it is present. May result in ineffective MAS (wasted effort) due to recipient lines being classified as QTL[-] and thus the QTL being introgressed, when in fact it is already present. |
| Utility | 100% | Marker can be used to track/introgress QTL across only a narrow range of germplasm. For most populations alternative markers are needed. |

### Applying quality metrics: Evaluation of existing marker systems Technical metrics: Indels vs. SSRs

Since technical metrics relate to the performance of a specific marker assay, they are by necessity empirical and may vary widely between markers even when these have the same biological properties and are run on the same platform. Indeed, wide variation was seen between markers for both call rate and clarity, even within trait-specific indel markers in specific QTL regions such as qDTY4.1 and qNa1L (Fig 5).

**Fig 5.**
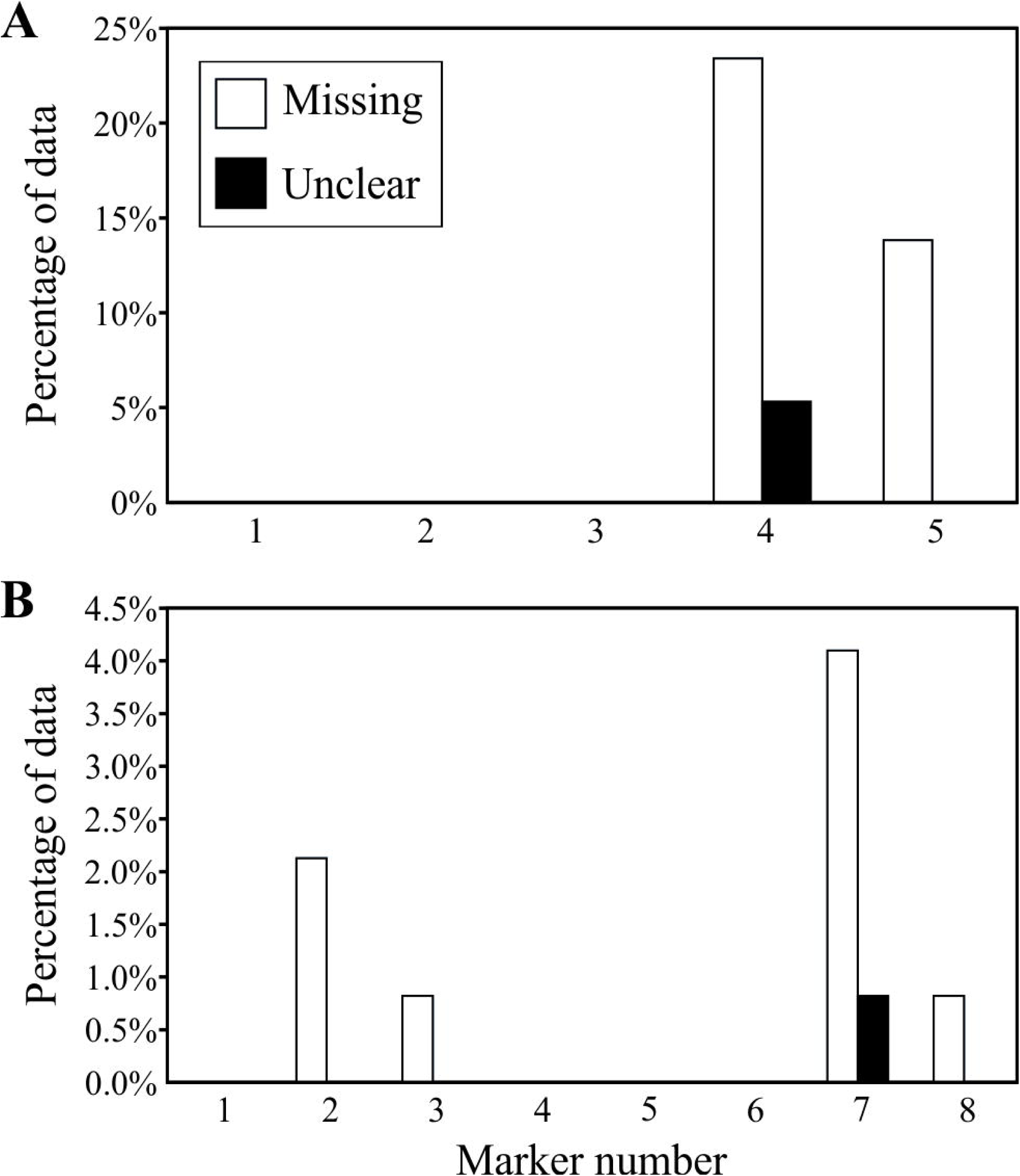
Technical metrics for candidate indel markers. Markers were evaluated within the qDTY4.1 (**A**) and qNa1L (**B**) QTL regions. Significant variation was seen for different markers in both QTL regions, showing some markers clearly performed better than others.

Technical metrics can be used to assess the relative performance of platforms as well as specific markers. The mean call rate and clarity were compared between a set of SSR and QTL-specific indel markers on a panel of 122 diverse cultivars (Fig 6). Trait-specific indel markers significantly out-performed SSR markers for clarity (P < 0.05). They also scored better than SSRs for missing data, though this was not significant (0.05 < P < 0.12), and may reflect a greater-than-average contribution from a few specific indels that scored very poorly, due largely to a few markers that covered genomic deletions in *qHTSF4* and *qDTY4.1*.

**Fig 6.**
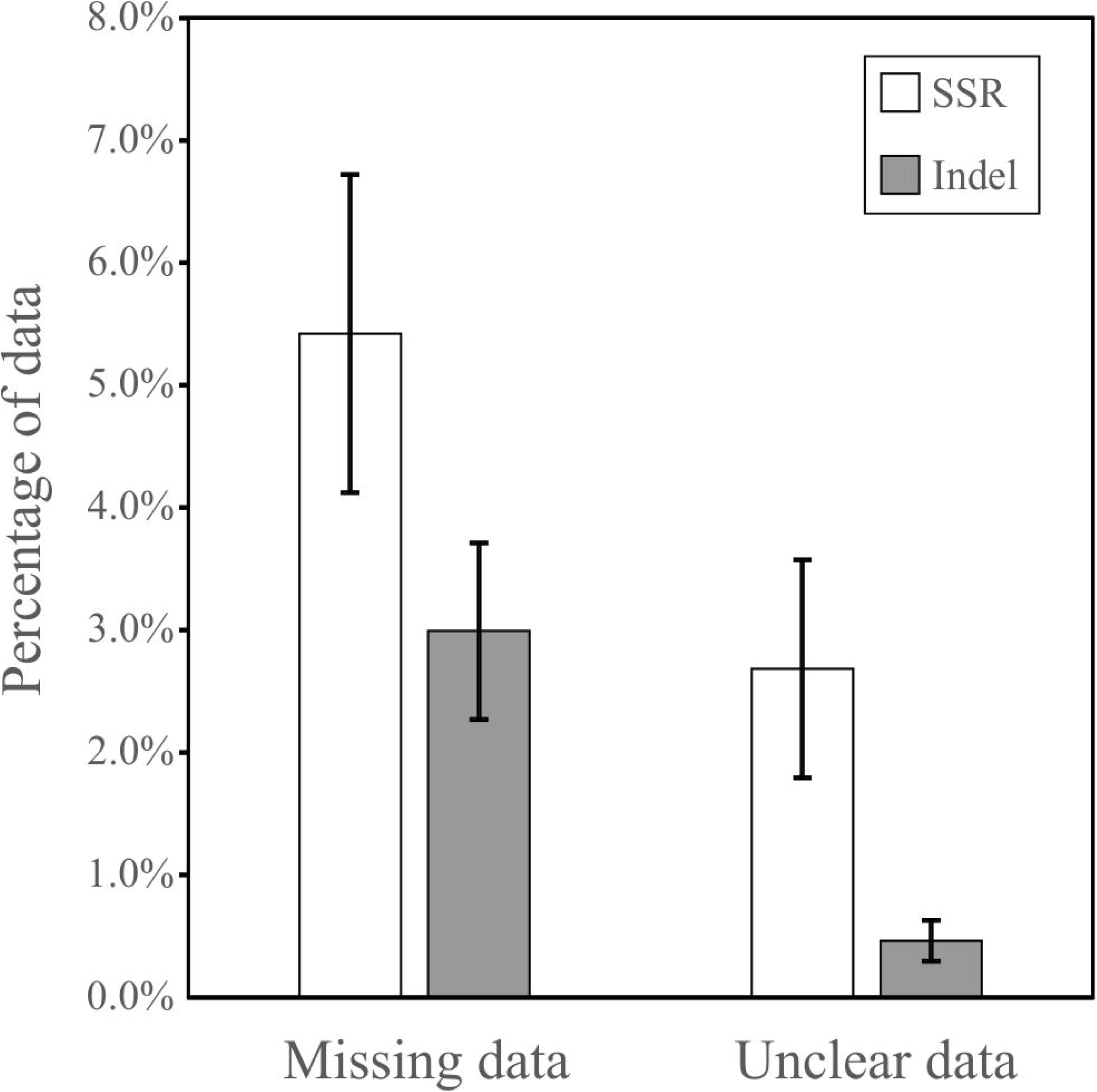
Comparison of performance of PAGE-based marker systems. The performance of SSRs was compared to that of trait-specific indels. Indels consistently out-performed SSRs, particularly in marker clarity but also in call rate.

### Biological metrics: Anonymous vs. Trait-specific markers

Working with whole-genome resequencing data it is evident that numerous polymorphisms can be easily identified between two varieties. However, these polymorphisms vary widely in their level of association with a target QTL. The level of association (FPR and FNR) for candidate markers throughout a salinity tolerance QTL in rice between 37 and 41Mb on the long arm of chromosome 1 shows wide variation, all through the QTL interval (Fig 7). This shows linkage with a QTL is not sufficient to give reliable selection. Secondly, none of the anonymous SNPs found on the Infinium chip within the QTL region (Fig 7a) scored perfectly on both the FPR and FNR, indicating they all suffer from errors in classifying known varieties. These SNPs have been filtered for those which show polymorphism between known donors and recipients, and corrected to minimise the false negative rate (correctly identifying as many donors as possible). In contrast, while QTL-specific indel and SNP markers (Figs 7b and c) also show variation in their association across the QTL, both classes have several markers achieving ideal scores (0%) on both metrics. Those markers scoring >0% on either the FPR or FNR may find niche applications in fine-mapping or specific populations, but those with perfect scores would make the most reliable marker system for breeding purposes. These metrics thus give information allowing the identification and design of optimal marker systems, even for quite large QTL regions.

**Fig 7.**
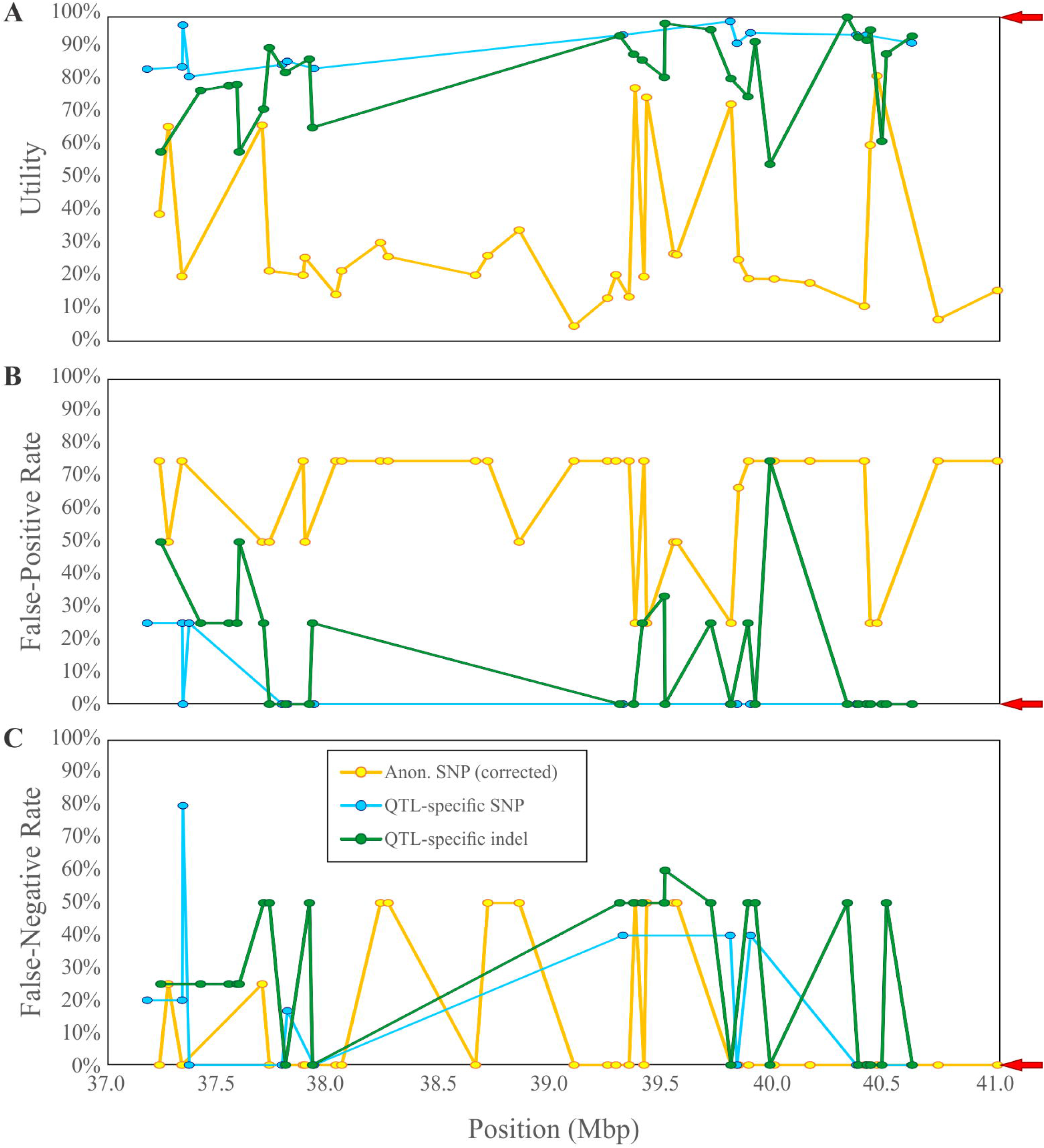
Use of marker quality metrics to determine optimal markers for a QTL. Comparison of quality metrics for different markers within a QTL region for salinity tolerance, qNa1L, between 37 and 41Mb on the long arm of chromosome 1. Multiple markers from the Illumina Infinium chip (Anonymous SNPs; favourable allele corrected to minimise FNR), QTL-specific SNPs and QTL-specific indels were assessed for their utility (**A**), false-positive rate (**B**) and false-negative rate (**C**) across IRRI germplasm. Anonymous SNPs typically scored poorly on FPR, FNR and Utility, and none scored well on all metrics. QTL-specific SNP and indel markers typically showed low or perfect scores on the FPR, and several markers scored 0% (no misclassified entries) on both FPR and FNR metrics. In addition QTL-specific markers scored far better for utility, indicating wider applicability in breeding. Arrows to the right indicate ideal target values for a new marker.

Improvements in the false positive and negative rates with QTL-specific markers are also seen for other QTL targets. Mean values from 42 QTLs for a range of traits including stress tolerance, grain quality, disease resistance etc. show that anonymous SNPs derived from the 6k diversity set consistently under-perform (Fig 8). Re-assignation of favourable and unfavourable alleles to minimise the false negative rate improves that metric to a level equivalent to those seen for the QTL-specific markers, but no better (P > 0.05). Optimising the FNR also improved the FPR, but not to a level equivalent to QTL-specific markers, so anonymous markers still performed worse on average (P < 0.0001). This means these anonymous markers, even with “corrected” assignations of favourable allele, have no relative benefit in detecting presence of the QTL, but do have a significant penalty in their FPR, incorrectly classifying lines as QTL[+].

**Fig 8.**
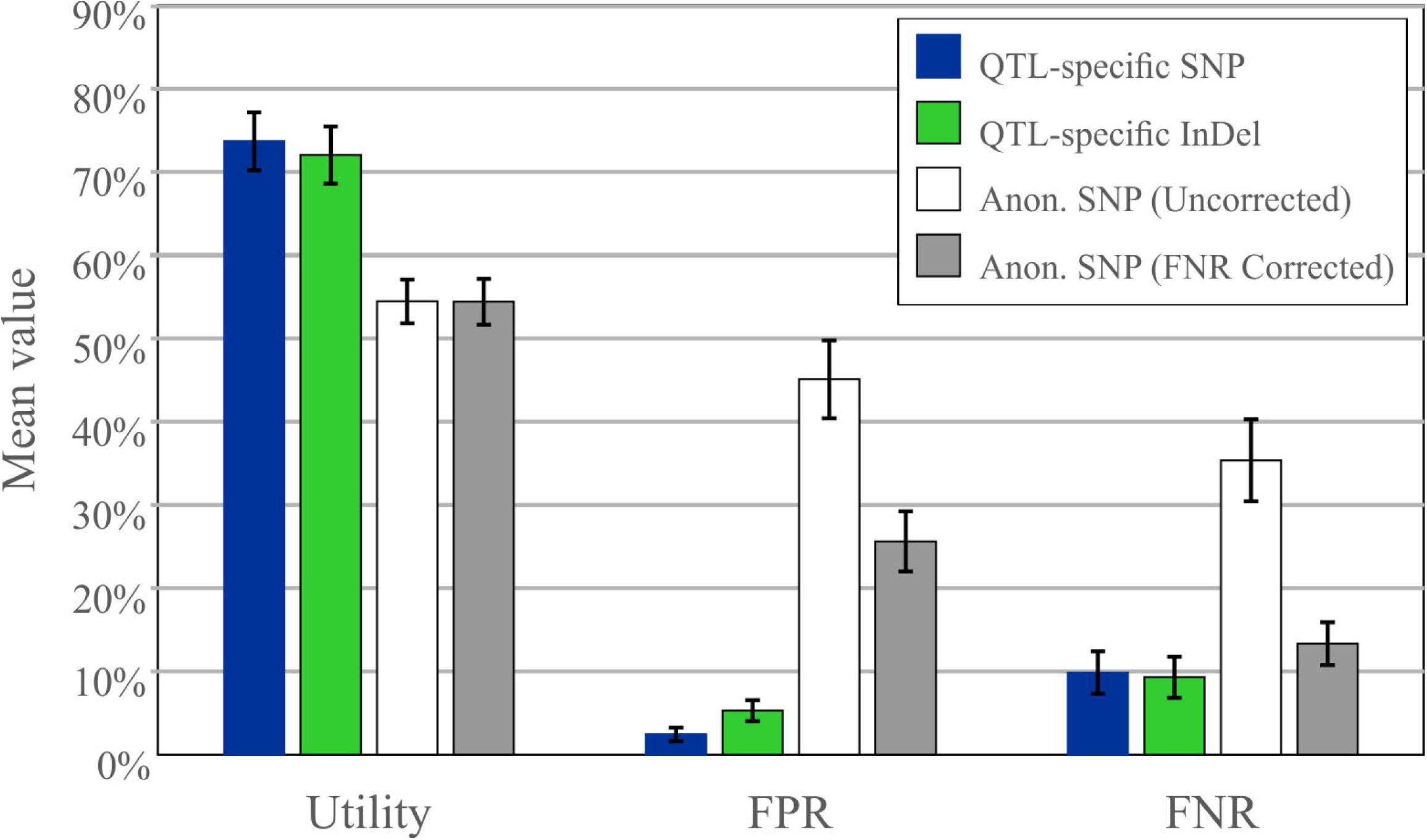
Comparison of mean accuracy metrics for diversity SNPs and QTL-specific SNP and indel markers. Anonymous SNP markers initially have very low scores on both the FPR and FNR. Correcting the assignation of favourable and unfavourable alleles to minimise the FNR improves that score to equivalent levels with the QTL-specific markers, but the FPR remains poor and no benefit is seen for the utility.

### Breeding metrics: Utility

Anonymous SNPs also showed lower average utility values (Fig 8), i.e. the designated favourable allele is present at higher frequencies in elite germplasm, and so the marker is less useful for introgressing a given QTL into a range of elite material. The utility metric is especially useful in the case of diagnostic markers, as it then indicates directly the proportion of elite material that a QTL may improve. Utility values for a range of QTL controlling various yield, grain quality, disease resistance and stress tolerance traits show a wide range in variation (Fig 9). These range from less than 20% for *LTG1, qSCT1* and *SCM2*, which appear to be fixed in nearly all *indica* elite material, to 100% for many disease resistance QTL. The latter observation is surprising considering the substantial selective pressure exerted on disease resistance in most breeding programs, and further work to determine its cause seems warranted.

**Fig 9.**
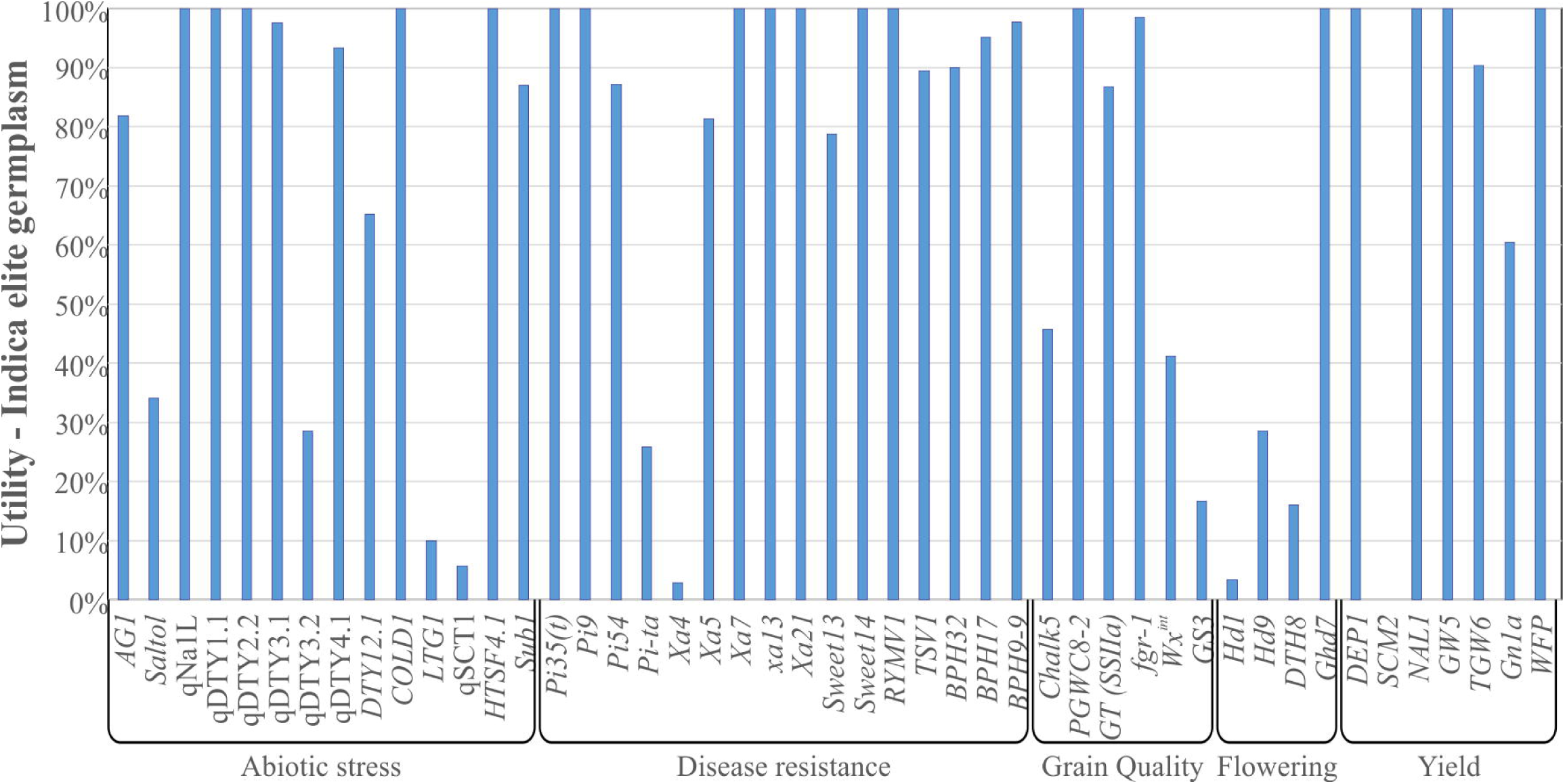
Variation in utility between various QTL for agronomic traits in *indica* breeding germplasm. QTL were selected that have diagnostic markers or markers scoring 0% on both FPR and FNR (and thus could be accurately scored). Wide variation in QTL utilities were seen, from near-fixation (utility ∼0%) to absent (utility 100%), but most were rare or absent.

## Discussion

Since the 1980s with the advent of SSR markers, it has become almost a mantra that the ideal marker should be highly polymorphic. This is certainly a useful feature for certain applications such as in bi-parental mapping, where high polymorphic information contents (PIC) increase the chances a given marker will be polymorphic between random parents. However, marker-assisted selection places different demands on the markers – the number of alleles displayed by the marker is not relevant, rather the ability to unambiguously discriminate between all donor and recipient material becomes critical.

Surprisingly, there is a dearth of literature on designing reliable markers and almost no criteria for judging what makes a marker “good” or “bad”. Most MAS programs use markers identified in QTL mapping populations – typically SSRs, applying them to other genetic backgrounds, and even attempting to use them to determine the presence of a QTL in diverse germplasm panels. These applications require very stringent false positive and false negative rates, but few examples exist where some validation of these false positive and negative rates has been conducted. Bernardo *et al.* [20] examined the reliability of SSR markers in selecting for stem rust resistance in wheat. Association of the markers with donor and recipient germplasm was analysed, but not quantified as a metric; association was not always good, for example markers for *Sr32* targeted the recipient allele not the (derived/wild) donor allele, and so run the risk of classifying some lines without the QTL as positive. Similarly, markers for other resistance genes variously failed to distinguish some or all known recipients from known donors (poor FPR; many SSRs had this difficulty), while others showed poor separation of donor and recipient alleles, produced major stutter bands, or produced non-target amplicons (poor clarity, figures 1 and 3 in their paper). In another strategy, Mohammadi-Nejad *et al.* [15] performed allele mining of the *Saltol* QTL using SSRs. Of particular note is their table 3, which lists the SSR haplotype and varieties which possess this haplotype. This can be related to alleles of the *HKT1;5* gene [23], which is causal for this QTL [29]. This reveals the SSR markers confounded (failed to distinguish) lines that had different alleles, such as IR64 and Kala Rata. Likewise, the converse was even more common: IR64 / IR29, and Pokkali / Kala Rata / Sadri were placed in separate haplotypes while sharing the same allele at the causal gene. Thus again the SSR genotype, and even haplotype, was not sufficient to reliably classify donor and recipient material for the QTL. In a counter-example, Tian *et al.* [30] designed indel markers for the allelic major rice bast resistance gene *Pi2*/*Pi9* based on sequence comparisons of parental varieties. The FPR and FNR were not quantified, but the inclusion of multiple reference alleles presumably helped in the design of markers highly specific to the favourable alleles. This marker was then able to demonstrate near-zero occurrence of this gene in a set of Chinese breeding germplasm, where previous marker sets were prone to false positives (see [31]) – incidentally indicating a potentially very high utility, though this was not articulated as a metric. A similar effort was conducted by Scheuermann and Jia [32] using a different approach. The latter marker from Scheuermann and Jia [32] should have the same FPR and FNR as that designed by Tian *et al.* [30], but suffered very low apparent call rates and clarity. In neither case were any metrics similar to FPR, FNR, Utility and Clarity quantified by either group, despite their datasets being sufficient to do so. Had these metrics been quantified, they could easily demonstrate which marker system is better and how reliably these markers could be used in other breeding programs, thereby greatly enhancing the impact of this work.

These examples show the need for a better system for describing the association of a marker with its target QTL. The fourteen metrics described in Table 1 are a substantial step towards providing such a description. Association of a marker with its target QTL is captured by a range of biological metrics rooted in the preceding discussion on the evolution of markers. Additional metrics describe parameters relating to reliability of the genotyping information, and the applicability of a marker in specific breeding situations. The preceding considerations have shown how the ideal marker – one which reliably identifies all donor and recipient germplasm – is based on the same polymorphism as gives rise to the QTL phenotype. Such a marker can be called diagnostic, and requires the identification of the gene *and the mutation* giving rise to a QTL – something that is very rarely done, even in rice. While ideal, this is difficult and time consuming. Alternative, flanking markers can still accurately classify observed alleles provided they arose at similar times and in the same lineage as the causative mutation (i.e. the derived marker allele matches the derived QTL allele; Fig 3). Again, the metrics in Table 1 provide a means to evaluate and validate candidate markers before committing to design and implementation (very important for expensive SNP systems) as well as assess performance after implementation. Validating existing marker systems with these metrics illustrates several points. First and foremost, QTL-specific marker systems consistently out-perform both older SSR and new anonymous SNP systems in most of these metrics, but most notably in the accuracy metrics FPR and FNR. For many QTL, no anonymous markers showed the required level of association with the target QTL. Thus QTL-specific markers will give consistently more reliable results in selection. In addition, accurate markers (scoring 0% on both FPR and FNR) can be used to determine the proportion of a breeding panel that may benefit from that QTL – the utility. Utility values vary widely between QTL (Fig 9), which reflects a complex interplay of the QTL’s origin and the artificial and natural selective pressures it has been subjected to in breeding programs. For example, *SCM2* is widely regarded as a candidate to reduce lodging, a major problem even in semi-dwarf rice. Unfortunately, however, the characterised donor allele of *SCM2* from Habataki [33] appears identical to that already found in the vast majority of *indica* breeding germplasm. This means the donor allele is already present in most or all improved *indica* material, and therefore the Utility of *SCM2* (and accurate markers for this gene) is very low in most *indica* breeding programs. By contrast many of the disease resistance loci have high utility, despite being under strong selective pressure in breeding programs.

Secondly, there is no inherent advantage of SNP genotyping platforms in terms of selection accuracy. Existing anonymous SNP marker sets such as the 6k Infinium chip were designed to maximise the probability of polymorphism (discriminatory power) between randomly-chosen varieties [6]. This makes them ideal for population genetics and as a fixed panel of markers to genotype any random set of parents and progeny. Ironically though, this means they have the least power to track specific QTL, and indeed they perform rather poorly in accuracy metrics overall. By contrast, QTL-specific markers, whether high-throughput SNP or low-throughput indel, perform quite well on accuracy metrics – and it is certainly possible to use these metrics to identify both SNP and indel markers with “perfect” associations with their target QTL (Fig 7). Therefore, the choice of marker *platform* has less to do with selection accuracy than with the expected sample throughput. However, the choice of best *marker* will be based on the level of association with the target QTL. Thirdly the application of the metrics in evaluating individual markers is easily demonstrated (Figs 5 and 7). Biological and breeding metrics are mostly useful in distinguishing between candidate markers prior to committing time and resources to implementing these on a particular genotyping platform; these are about choosing the optimal target polymorphisms. Yet although a candidate marker may have perfect biological and breeding metrics, a given assay for that polymorphism may perform very poorly on its technical metrics (call rate and clarity), such as seen for *qDTY4.1* (Fig 5). For PCR-based systems such as SSRs, indels and some SNP technologies, much of this variation in the technical metrics is due to inherent issues with primer efficiency. However, while SNP assays also utilise oligonucleotides as either probe or primer sequences, it is unfortunately rare to see “validation” of the performance of new markers on SNP platforms due to the expense of an assay. In addition, SNP assays very rarely return the raw data, instead reporting a digested summary –the actual SNP call–thus glossing over such factors as whether the clustering of fluorescence intensities was unambiguous. It seems advisable going forward to implement some form of technical and biological replication when validating a new assay to determine its clarity.

Allied to this discussion on technical metrics, it is evident that a given polymorphism may be interrogated by multiple different “markers”. For example, a SNP could be targeted by one of several gel-based assay systems, any of the many SNP platforms, or an amplicon/sequencing approach. All are based on the same polymorphism, but as each platform has its own design quirks and even a single platform may use alternate primer pairs for amplification, different instances of a marker could have very different scores on technical metrics, despite targeting the same polymorphism. Therefore, information on the version/instance of a marker under consideration is needed to distinguish between alternate forms and platforms targeting the same polymorphism. These quality metrics thus provide a good framework for assessing the accuracy and reliability of any specific marker. This information can be used to evaluate existing markers, design better markers, and even to compare performance of different marker types and platforms. Nevertheless, they are by no means complete or perfect. Two issues worth highlighting concern the Clarity metric and estimation of the FPR and FNR metrics.

The Clarity metric is currently slightly ambiguous; it could refer to either how clearly/reliably genotyping data (bands on a gel, fluorescence signal clusters on a SNP platform, or other measures) can distinguish between the allelic states of the marker. It could also refer to how often duplicate samples cluster together – repeatability. This is of course closely related to the former situation, but is also subtly different. For the sake of simplicity these are not distinguished here, but further discussion on whether Clarity as described here should be broken down into two metrics – clarity and repeatability – seems warranted.

Of the metrics presented, the accuracy metrics FPR and FNR are arguably the most important in distinguishing between candidate markers. A common objection and difficulty in the assessment of these is that they require a large number of case-[+] and case-[-] data points to accurately estimate, whereas in most cases the number of defined donor and recipient lines is limited. This is absolutely true – and should inform how QTL mapping and validation efforts are undertaken – but is not a justification to reject the importance of these metrics. On one hand, some estimate is better than none, and it should be recognised that all datasets are inadequate, to some extent. It is thus better to report the statistics, together with data on how accurate these might be – such as the total number of defined donor and recipient lines available, where a greater number of both implies greater accuracy in estimation. On the other hand, estimates of FPR and FNR based on inadequate (small) datasets actually *inflate* their values. In the extreme case of a single known donor and recipient line, any marker polymorphic between these parents will then score 0% on both metrics. An inadequate dataset thus does not lead to a rejection of markers, but rather the opposite: the power to distinguish between them is limited, and so any feature that shows polymorphism will be accepted. In this situation an educated guess as to whether the favourable or unfavourable allele is likely to be derived (i.e. the derived QTL allele). Markers targeting this (i.e. the marker target allele is the same as the derived QTL allele) are then effectively making the assumption that the derived QTL allele is rare in the overall allelic diversity, thus deliberately biasing the error towards false-negatives and away from false positives – maximising the probability of a good FPR at the potential penalty of FNR – and thus biasing risk away from advancing unfavourable genotypes at the penalty of increasing risk of discarding favourable ones.

In summary, the metrics proposed in Table 1 quantify all significant parameters describing a marker’s behaviour when assayed, the level and type of association it displays with its target QTL (both species-wide and in specified breeding panels), and its distribution within a breeding panel. These metrics give a fast, comprehensive and objective means to discriminate between and evaluate alternative markers (e.g. Fig 7), allowing an optimal marker system to be designed. In addition, by including such “housekeeping” metrics as the favourable and unfavourable alleles, it is possible to automate these calculations, providing the possibility to scan genomic datasets for optimal SNP markers programmatically, greatly simplifying the deployment of QTL in breeding. It also allows the development of a marker database with live updating of metrics as new data is added, enabling continual refinement of marker systems. The advantages of adopting a set of metrics are manifold, and it is hoped that the proposed system will assist in developing a new generation of reliable marker systems to improve the efficiency of plant breeding.

## Acknowledgements

The authors wish to thank Irish Bagsic, Chenie Zamora and Katreena Titong for technical assistance in various aspects of this work.

## Supporting information

**S1 Table. List of indel and SSR markers assessed for technical performance.**

**S2 Table. List of varieties examined for technical performance evaluation.**

**S3 Table. List of varieties examined in calculating biological accuracy and breeding metrics.**

**S4 Table. List of marker positions interrogated for assessing biological accuracy and breeding metrics.**

**S5 Table. List of QTL examined, with start and end positions.**

